# VData: Temporally annotated data manipulation and storage

**DOI:** 10.1101/2023.08.29.555297

**Authors:** Matteo Bouvier, Arnaud Bonnaffoux

## Abstract

**Background:** Recent advances in both single-cell sequencing technologies and gene expression simulation algorithms have led to the production of increasingly large datasets. Larger datasets (tens or hundreds of Gigabytes) can no longer fit on regular computers’ RAM and thus pose important challenges for storage and manipulation. Existing solutions offer partial solutions but do not explicitly handle the temporal dimension of simulated data and still require large amounts of RAM to run.

**Results:** VData is a Python extension to the widely used AnnData format that solves these issues by extending 2D dataframes to 3 dimensions (cells, genes and time). VData is built on top of Ch5mpy, a custom built Python library for easily working with hdf5 files and which allows to reduce the memory footprint to the minimum.

**Conclusions:** VData allows to store and manipulate very large datasets of (empirical or simulated) time-stamped data. Since it follows the original Ann-Data format, it is compatible with the scverse tools and AnnData users will find it easy to use.

## Background

Since their introduction in [1], single-cell sequencing protocols have revealed with unprecendented details the highly dynamic and heterogeneous nature of gene expression. There now exists a wide variety of such protocols (10x Genomics [2], SMART-seq [3], MARS-seq [4], Split-seq [5]) which were leveraged by the systems biology community to gain insights into how genes are expressed and regulated inside cells. As a result, vast amounts of single-cell expression datasets have been produced, with an ever increasing number of cells and genes being captured in a single experiment. It has now become common to analyze datasets of tens or hundreds of thousands of single cells characterized by the expression levels of tens of thousands of genes [6]. This is even truer in multiomics studies, in which multiple genetic parameters are measured in the same unique cell in parallel, thereby increasing again the size of the datasets (scMT-seq [7], SHARE-seq [8], FISH-Flow [9]).

Experiments are not the only source of gene expression datasets. Synthetic datasets are now beeing produced by a variety of simulation algorithms, used to test our understanding of gene expression and regulation mechanisms or to predict cellular cell fate (dyngen [10], Splatter [11], scDesign2 [12], SNCS [13], WASABI [14], CAR-DAMOM [15]). Although similar to experimental datasets, synthetic data displays unique characteristics since 1) it is often non sparse (there are no technical dropouts and simulations are approximate), 2) there is potentially no limit to the number of cells or genes for which to record data and 3) simulations allow to observe the same cell at multiple timepoints (although some experimental protocols now also allow this, such as Live-seq [16]).

Experimental datasets, and synthetic ones even more so, have become extremely large, to the point that they now frequently reach several Gigabytes. As data analysis progresses, the total size of the datasets still increases as copies are made or new matrices are created. This poses important computational challenges that need to be addressed to ensure that current computers, with limited memory size, are still able to work on such large datasets.

Algorithms dedicated to single-cell dataset storage and manipulation have been proposed (AnnData [17] in Python, SingleCellExperiment [18] in R) but do not specifically focus on working with large datasets (only the principal matrix in AnnData objects – the X layer – can be read from an on-disk file one small chunk at a time) or with synthetic data (there is no explicit handling of the time dimension, preventing the exact same cell to be found multiple times).

Here we present *VData*, a solution for storing and manipulating single cell datasets that extends the widely used AnnData format and is designed with synthetic data in mind. Most notably, it adds a third ‘time’ dimension beyond the usual ‘cell’ and ‘gene’ axes to support time stamped longitudinal data and heavily focuses on low memory footprint to allow fast and efficient handling of large datasets of tens of Gigabytes even on regular laptops. VData is available as a Python package at https://github.com/Vidium/vdata.

## Implementation

### General VData structure

The general VData object structure matches as closely as possible the one of AnnData objects (see Fig 1). As VData introduces an additional time dimension however, a supplementary ‘timepoints’ dataframe exists to store data describing individual time-points. Also, ‘layers’, ‘obs’ and ‘obsm’ are now 3D dataframes (*cells × genes × time*) instead of 2D (*cells × genes*).

**Fig. 1.**
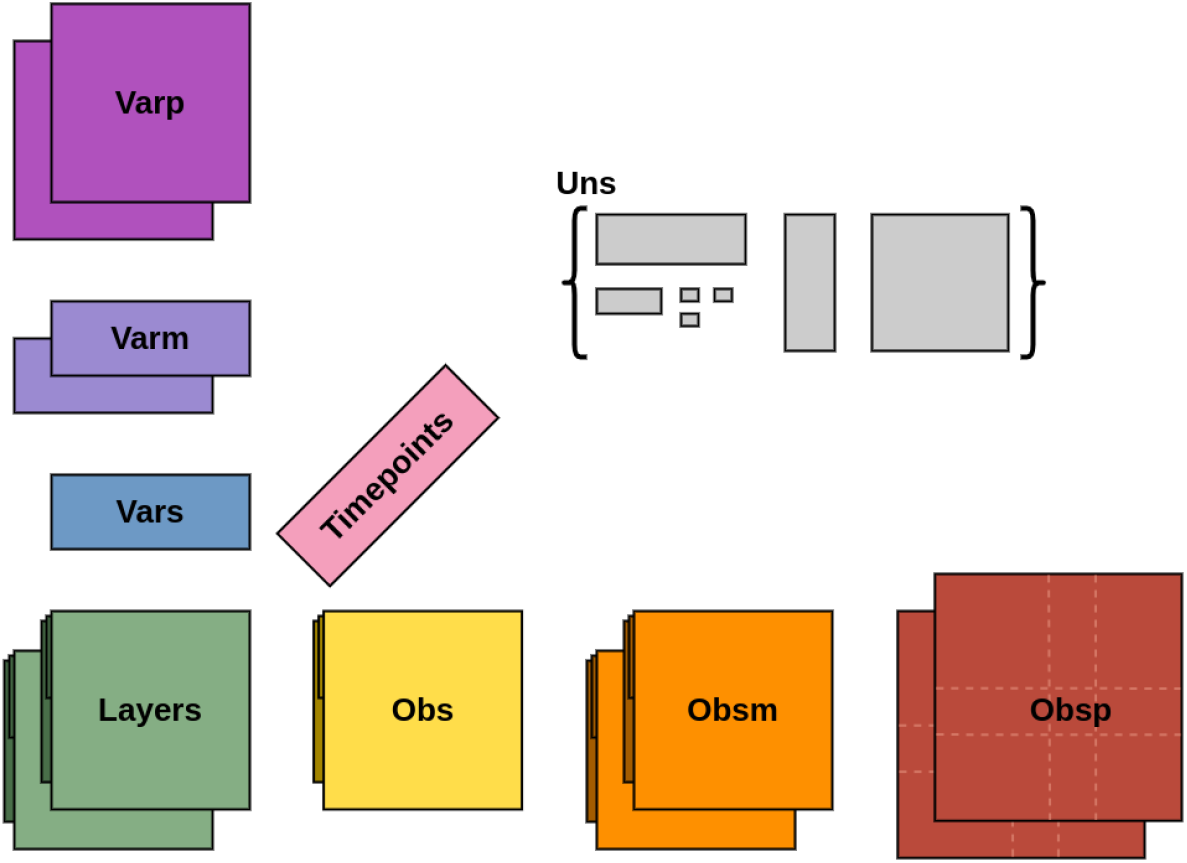
Structure of a VData object. As in AnnData objects, the bulk of the data is stored in multiple matrices called ‘layers’. The data is described using ‘obs’ and ‘var’ dataframes for providing details about individual cells and genes respectively. Multi-dimensional or pair-wise data can be stored in ‘obsm’, ‘obsp’, ‘varm’ and ‘varp’ mappings (see [17] for details). Additionally, data independent from cells and genes can be stored in the ‘uns’ mapping.

### VData introduces temporally annotated dataframes

To easily attribute data to a particular timepoint or to retrieve it, we needed to expand classical pandas DataFrames [19] to three dimensions. For that, we created TemporalDataFrames, providing usual features of pandas DataFrames and subsetting mechanisms with an extra time dimension. Under the hood, TemporalDataFrames work on 2-dimensional datasets but store an extra hidden column for annotating rows with the timepoint they belong to. The dataset is only reorganized into 3 dimensions for display. Obtaining a subset of a TemporalDataFrame is done with the usual square brackets indexing mechanism, where the first index selects one or more timepoints, the second index selects rows (cells) and the third selects columns (genes) (see **S1 Listing** for an example). For interoperability with pandas, TemporalDataFrames can be created from pandas DataFrames or can be converted to them.

### VData can read from on-disk files to reduce the memory footprint

Once a VData object has been created, it can be stored as an on-disk file using the hdf5 file format [20] (see Fig 2) similarly to how it can be done in AnnData. Opening a VData from an hdf5 file is refered to as “backing” the object. When a VData object is backed – and as opposed to how it is done in AnnData – **all** of its contents are accessed in small chunks of configurable size only when required. This includes ‘layers’, ‘obs’, ‘var’, ‘timepoints’ but also all data in the ‘uns’ dictionary. This means that no object needlessly uses computer memory, except for the one currently being read or written to.

**Fig. 2.**
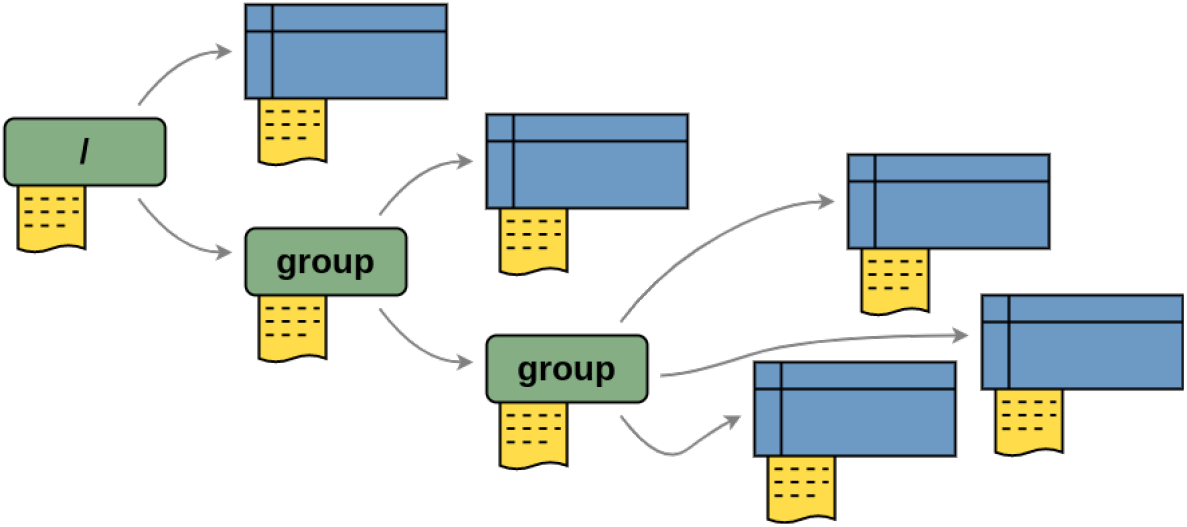
HDF5 file format. HDF5 files contain a tree of containers called *groups* (green boxes) and *datasets* that contain data of any dimension (blue boxes). The root of any HDF5 file is the “/” group. Groups can contain any number of sub-groups or of datasets. Metadata, called *attributes*, can be attached to both groups and datasets to add information (in yellow).

To allow data of many different types to be backed, we wrote an other small Python library called *Ch5mpy*.

### VData is built on top of Ch5mpy for reading from hdf5 files

Ch5mpy (pronounced “champy”) is a small Python library built on top of the h5py library [21] that introduces user-friendly wrappers to handle objects and data stored in hdf5 files as more common Python objects. Most notably, Ch5mpy allows to :

- explore an hdf5 file as if it were a Python dictionary, only loading in memory the groups or datasets names. The actual dataset will be loaded only when requested by the user.
- interact with datasets as if they were numpy arrays [22]. This includes subsetting, reshaping, data-type casting and applying numpy operations while ensuring the memory usage stays below a configurable threshold.

It also defines a simple API for writing and reading any object to an hdf5 file as a collection of datasets and in some cases will guess on its own how to write simple objects to the file. Ch5mpy is available as a standalone Python package at https://github.com/Vidium/ch5mpy and can be installed from PyPI (https://pypi.org/project/ch5mpy/).

## Results

### Single cells can be tracked at multiple timepoints

To handle time-stamped data easily, VData accepts indices with repeating values so that the exact same cell name can be found at all timepoints. This avoids having to modify the index (such as appending the timepoint id to the cell names) to create valid objects. An example of TemporalDataFrame generation with repeated cell names is given in **S2 Listing**.

### VData opens large datasets with low memory footprint

To demonstrate VData’s ability to load very large datasets, we built test VData and AnnData objects, each with 4 layers of random values with increasing numbers of genes and cells :

- 1000 cells *×* 2000 genes (62.8MB)
- 10000 cells *×* 10000 genes (3.1GB)
- 50000 cells *×* 10000 genes (14.7GB)
- 50000 cells *×* 50000 genes (75.1GB)
- 100000 cells *×* 50000 genes (149.8GB)

As shown in Fig 3.a, since VData only creates references to on-disk data objects and accesses it only when necessary, it was able to load all datasets while maintaining the memory footprint to less than 100MB. This allowed VData to read datasets larger than the available amount of RAM on our computer and in less than 0.1 seconds. On the other hand, AnnData required large amounts of RAM and was slow to load large datasets. This was expected since AnnData usually relies on sparse matrices to lower the memory footprint and load time, which is not the case in this test setup and often not true with simulated data.

**Fig. 3.**
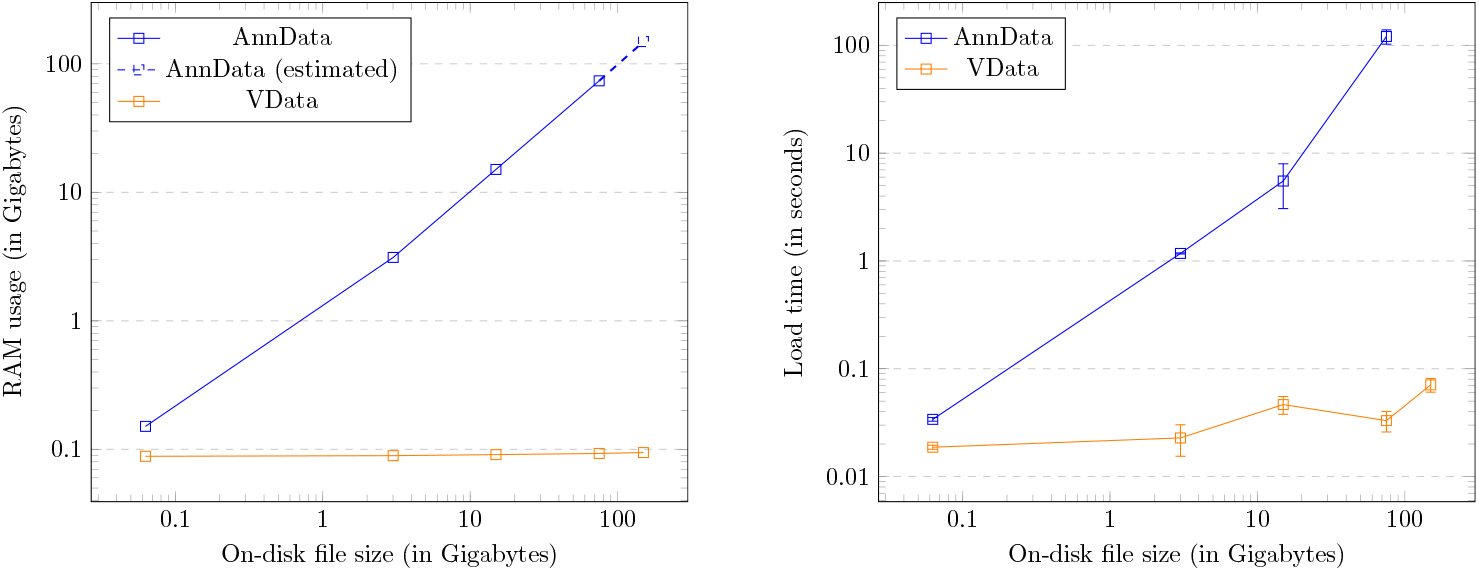
Dataset loading performance. **Left** Memory usage when loading datasets of increasing sizes from on-disk files. AnnData performance is recorded in blue, VData in orange. Final performance (with the 149GB file) could not be obtained on our computer and is only estimated. **Right** Loading speed when reading the datasets. AnnData performance is recorded in blue and VData in orange. Values reported are the mean of 50 read times and standard deviations.

### VData objects are compatible with the scverse ecosystem

We developped VData as an extension of Anndata objects, following its structure as closely as possible to make it easy for users to transitition to VData. Creating a VData object from an existing Anndata simply requires the user to initialize the VData from the Anndata as : ‘VData(existing anndata)’ or by calling the ‘read from anndata(“path/to/anndata”)’ method to create a VData from a saved AnnData file. If needed, VData objects can be converted back to Anndata by calling the ‘to anndata()’ method.

To allow VData objects to be used directly by tools of the scverse [23] without first converting VData objects to AnnData and thus giving up on the efficient memory usage, VData objects can be wrapped into an ‘AnnDataProxy’. AnnDataProxies inherit from AnnData and parse AnnData operations or method calls to produce valid VData operations. With this, we successfully ran the scvelo [24] ‘Getting Started’ tutorial, passing a VData wrapped in an AnnDataProxy to scvelo’s functions with no error.

## Conclusions

VData is an extension to AnnData written from scratch in Python. While the base structure remains the same, the supplementary time axis allows to easily deal with time-stamped data, often generated by simulation algorithms. When designing VData, focus was put on keeping the memory footprint low and on ease of integration with existing tools. For that, VData uses the Ch5mpy Python library (also made freely available) for backing all data on an hdf5 file and implements a wrapper that makes VData objects compatible with tools working on AnnData objects. This makes it possible to apply, on regular computers, widely used bioinformatics tools to datasets of sizes that would previously require enourmous amounts of RAM.

## Availability and requirements

**Project name:** VData **Project home page:** https://github.com/Vidium/vdata **Operating system:** Linux, Windows, OS X **Programming language:** Python3 **Other requirements:** NumPy, SciPy, Pandas and AnnData **License:** CeCILL-B **Any restrictions to use by non academics:** None

### List of abbreviations

API: ApplicationProgramming Interface
hdf5: Hierarchical Data Format version
RAM: Random Access Memory
MB: MegaByte
GB: GigaByte

## Declarations

### Ethics approval and consent to participate

Not applicable

### Consent for publication

Not applicable

### Availability of data and materials

VData and Ch5mpy are available as Python libraries. They can be downloaded from github at https://github.com/Vidium/vdata and https://github.com/Vidium/ch5mpy respectively. Libraries can also be installed with pip by running ‘pip install vdata’ which installs both VData and Ch5mpy, or by runninh ‘pip install ch5mpy’ which installs Ch5mpy alone.

### Authors’ contributions

MB designed and developed VData and Ch5mpy. MB and AB drafted the manuscript. All authors read and approved the final manuscript.

## Acknowledgements

We would like to thank Elsa Guillot, Quentin Fort, Paul Simon and Olivier Gandrillon for their thorough testing, for suggesting improvements and features and in general for their precious feedback.

