## Supplementary material for "VData: Temporally annotated data manipulation and storage"

```python
import vdata as vd
import numpy as np

t = vd.TemporalDataFrame(
    np.random.rand(30, 5),
    timepoints=np.repeat([0, 1, 2], 10),
    index=[f'c{i}' for i in range(1, 31)],
    columns=['g1', 'g2', 'g3', 'g4', 'g5']
)

# >>> print(t)
# TemporalDataFrame No_Name
# Time point : 0.0 hours
# ----------------------------------------
#       Time-point           g1        g2        g3        g4        g5
# c1          0.0h  |  0.391084  0.813595  0.141663  0.841863  0.398797
# c2          0.0h  |  0.622206  0.362086  0.676001  0.485081  0.468641
# c3          0.0h  |  0.611089  0.931794  0.736403  0.975381  0.302012
# c4          0.0h  |  0.720324  0.431909  0.639728  0.432422  0.698777
# c5          0.0h  |  0.985004  0.085789  0.755214  0.336054  0.419931
# [10 rows x 5 columns]
#
# Time point : 1.0 hour
# ----------------------------------------
#        Time-point           g1        g2        g3        g4        g5
# c11          1.0h  |  0.602622  0.870247   0.24695  0.819429  0.505861
# c12          1.0h  |  0.191022  0.210838  0.146375  0.212947  0.219807
# c13          1.0h  |  0.812675  0.745807  0.535001  0.555916  0.162295
# c14          1.0h  |  0.988531  0.752326  0.294352  0.515573  0.970844
# c15          1.0h  |  0.618349  0.325882  0.337374  0.916267  0.442175
# [10 rows x 5 columns]
#
# Time point : 2.0 hours
# ----------------------------------------
#        Time-point           g1        g2        g3        g4        g5
# c21          2.0h  |  0.709283  0.370199  0.920588   0.32927  0.285588
# c22          2.0h  |  0.529812  0.254663  0.732573   0.98778  0.475708
# c23          2.0h  |  0.897081   0.19356   0.74047  0.109317  0.488225
# c24          2.0h  |  0.789474  0.927484  0.959109  0.926143    0.7216
# c25          2.0h  |  0.472701  0.040254  0.943085  0.602839  0.698247
# [10 rows x 5 columns]

# 1. Get a subset on timepoints by selecting only timepoint '0 hours'
t_0h = t['0h']

# print(t_0h)
# View of TemporalDataFrame No_Name
# Time point : 0.0 hours
# ----------------------------------------
#       Time-point           g1        g2        g3        g4        g5
# c1          0.0h  |  0.391084  0.813595  0.141663  0.841863  0.398797
# c2          0.0h  |  0.622206  0.362086  0.676001  0.485081  0.468641
# c3          0.0h  |  0.611089  0.931794  0.736403  0.975381  0.302012
```

```
# c4        0.0h  |  0.720324  0.431909  0.639728  0.432422  0.698777
# c5        0.0h  |  0.985004  0.085789  0.755214  0.336054  0.419931
# [10 rows x 5 columns]

# 2. Get a subset on cells by selecting all timepoints with ':' and by selecting cell named 'c1'
t_c1 = t[:, 'c1']

# >>> print(t_c1)
# View of TemporalDataFrame No_Name
# Time point : 0.0 hours
# ------------------------
#    Time-point          g1        g2        g3        g4        g5
# c1        0.0h  |  0.391084  0.813595  0.141663  0.841863  0.398797
# [1 rows x 5 columns]

# 3. Get a subset on genes by selecting all timepoints and cells and by selecting gene named 'g2'
t_g2 = t[:, :, 'g2']

# >>> print(t_g2)
# View of TemporalDataFrame No_Name
# Time point : 0.0 hours
# ------------------------
#    Time-point          g2
# c1        0.0h  |  0.813595
# c2        0.0h  |  0.362086
# c3        0.0h  |  0.931794
# c4        0.0h  |  0.431909
# c5        0.0h  |  0.085789
# [10 rows x 1 columns]
#
# Time point : 1.0 hour
# ------------------------
#     Time-point          g2
# c11        1.0h  |  0.870247
# c12        1.0h  |  0.210838
# c13        1.0h  |  0.745807
# c14        1.0h  |  0.752326
# c15        1.0h  |  0.325882
# [10 rows x 1 columns]
#
# Time point : 2.0 hours
# ------------------------
#     Time-point          g2
# c21        2.0h  |  0.370199
# c22        2.0h  |  0.254663
# c23        2.0h  |   0.19356
# c24        2.0h  |  0.927484
# c25        2.0h  |  0.040254
# [10 rows x 1 columns]
```

**S1 Listing Substetting TemporalDataFrames.** Code example of how to obtain subsets of TemporalDataFrames using the square brackets. Three elements can be used in the brackets: the first selects timepoints, the second selects rows (cells) and the third selects columns (genes). Final selection elements may not be given.

```python
import vdata as vd
import numpy as np

# build TemporalDataFrame with same index (cell names) at all timepoints
t = vd.TemporalDataFrame(
    np.random.rand(30, 5),
    timepoints=np.repeat([0, 1, 2], 10),
    index=vd.Index(['c1', 'c2', 'c3', 'c4', 'c5', 'c6', 'c7', 'c8', 'c9', 'c10'],
                   repeats=3),
    columns=['g1', 'g2', 'g3', 'g4', 'g5']
)

# >>> print(t)
# TemporalDataFrame No_Name
# Time point : 0.0 hours
# ----------------------
# Time-point              g1        g2        g3        g4        g5
# c1         0.0h  |  0.883432  0.557794  0.103124  0.167090  0.809125
# c2         0.0h  |  0.521437  0.290602  0.410404  0.011147  0.831306
# c3         0.0h  |  0.423411  0.336324  0.027670  0.815664  0.533868
# c4         0.0h  |  0.625875  0.432366  0.670179  0.297758  0.996317
# c5         0.0h  |  0.679658  0.266367  0.808400  0.415158  0.412760
# [10 rows x 5 columns]
#
# Time point : 1.0 hour
# ----------------------
# Time-point              g1        g2        g3        g4        g5
# c1         1.0h  |  0.165900  0.177450  0.312554  0.153819  0.493438
# c2         1.0h  |  0.789727  0.100523  0.585656  0.814834  0.781150
# c3         1.0h  |  0.842200  0.326500  0.408542  0.141354  0.379258
# c4         1.0h  |  0.774158  0.621878  0.943658  0.685533  0.248216
# c5         1.0h  |  0.721871  0.027862  0.340560  0.835714  0.091043
# [10 rows x 5 columns]
#
# Time point : 2.0 hours
# ----------------------
# Time-point              g1        g2        g3        g4        g5
# c1         2.0h  |  0.581677  0.432881  0.013921  0.016905  0.145276
# c2         2.0h  |  0.068850  0.959728  0.183135  0.042197  0.984561
# c3         2.0h  |  0.875083  0.681113  0.369171  0.875418  0.784618
# c4         2.0h  |  0.195309  0.535499  0.199410  0.651185  0.361248
# c5         2.0h  |  0.916437  0.010108  0.342436  0.106432  0.743724
# [10 rows x 5 columns]

# get subset of TemporalDataFrame, selecting cell 'c1' at all timepoints
t_c1 = t[:, 'c1']

# >>> print(t_c1)
# View of TemporalDataFrame No_Name
# Time point : 0.0 hours
# ----------------------
# Time-point              g1        g2        g3        g4        g5
# c1         0.0h  |  0.883432  0.557794  0.103124  0.167090  0.809125
# [1 rows x 5 columns]
#
```

```
# Time point : 1.0 hour
# ------------------------------
# Time-point             g1        g2        g3        g4        g5
# c1         1.0h   |   0.165900  0.177450  0.312554  0.153819  0.493438
# [1 rows x 5 columns]
#
# Time point : 2.0 hours
# -------------------------------
# Time-point             g1        g2        g3        g4        g5
# c1         2.0h   |   0.581677  0.432881  0.013921  0.016905  0.145276
# [1 rows x 5 columns]
```

**S2 Listing  Cells tracked at multiple timepoints in a TemporalDataFrame.** Code example of how to build TemporalDataFrames with cells tracked at multiple timepoints. This requires defining the cell names with vdata's Index with ‘repeats‘ set to the number of timepoints.
